# Efficient Protein Engineering via Integrated Language Models and Bayesian Optimization

**DOI:** 10.1101/2025.09.30.679490

**Authors:** Joshua Meehl, Prasad Siddavatam

## Abstract

This study investigates the application of advanced predictive models to reduce the cost and effort associated with protein engineering campaigns. We explore the use of protein language models (PLMs), a variant of large language models (LLMs), to predict functional performance from protein sequences. A common challenge in this domain is the scarcity of functional data. To address this, we examine zero-shot and few-shot learning methods. Another challenge is efficiently searching the vast fitness landscape for superior protein variants. We evaluate search methods, such as Bayesian optimization, to tackle this problem. The proposed methods are evaluated against a benchmark of 34 protein datasets containing sequences and their quantified functional values. Our findings demonstrate the potential of these advanced predictive models to streamline and accelerate the protein engineering process.

## 1 Introduction

Project Matterhorn explores how to efficiently select superior protein variants using computational methods. By leveraging state-of-the-art models and techniques, we aim to develop more efficient and effective approaches for modifying proteins to enhance their desired properties. These computational methods have the potential to significantly reduce the time and costs associated with traditional laboratory-based protein engineering techniques, while also producing better results. Ultimately, this work may contribute to the development of improved or novel enzymes.

The project addresses two primary challenges in computationally screening protein variants. The first is the *functional prediction problem*, which involves predicting the functional^*^ properties of a given protein variant when very little experimental data is available on those properties. The second is the *fitness landscape search problem*, which is how to efficiently explore the vast number of possible protein variants once property predictions are made, since exhaustively testing all variants is impractical.

This report provides the background and analysis of designing a computational screening pipeline for protein engineering. The rest of the Introduction section describes the pipeline, its components, and brief explanations and examples. The Literature Review section offers an overview of the state-of-the-art techniques in protein language models and Bayesian optimization that form the foundation of our approach. In the Results section, we present the outcomes and analysis of our screening pipeline, demonstrating its effectiveness in identifying promising protein variants. The Methods section delves into the technical details of the models, assumptions, and methodology employed in the pipeline, as well as the data used for its evaluation. Finally, the Discussion section summarizes the key conclusions drawn from our work and explores potential avenues for future research and development.

### 1.1 Computational Screening Pipeline

To address these two challenges, we designed a *computational screening pipeline*. This pipeline can evaluate millions of potential protein variants and select only the most promising candidates for experimental lab validation. Importantly, the pipeline can make reasonable initial predictions even without any prior functional data on the variants. The computational pipeline is coupled with a laboratory workflow, where the selected variants are physically cloned, expressed as proteins, and tested to obtain functional data. This experimental data is then fed back into the pipeline to improve the accuracy of predictions for the next experimental cycle of variant selection and screening As a hypothetical example, consider an engineering campaign with the goal of improving the activity of a wild-type (WT) enzyme^*^ by a factor of 2. The campaign is budgeted for 10 experimental cycles, with 100 protein variants tested in each cycle. In this situation, the computational screening pipeline would first make initial predictions of activity without any existing functional data on the variants. The pipeline then explores mutations to the WT enzyme sequence and predicts how those mutations will impact performance. The relationship between the sequence changes and their effects on performance is known as the *fitness landscape*. This workflow is shown in Figure 1.

**Figure 1.**
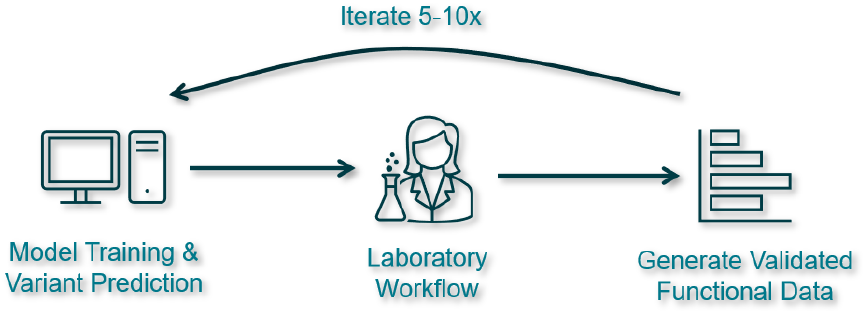
A diagram of the proposed workflow. The workflow starts with model training and computational variant screening. Selected variant sequences are provided to the laboratory team to clone and express the physical proteins. The functional values of the protein variants are then quantified and used to retrain the models. A campaign may have 5 - 10 experimental cycles.

Once this vast fitness landscape has been sufficiently explored computationally with millions of variants, a small subset of 100 promising variants is selected for actual laboratory testing. Some of these may fail to express properly or lack any enzymatic activity at all. Others may exhibit activity similar to the WT, while a small fraction could potentially show much higher activity. The experimental data from this first lab cycle is then fed back into the pipeline to improve its predictive models and better map the fitness landscape, including identifying both non-functional “holes” as well as potential peaks of high activity. These initial 100 data points allow the models to be fine-tuned, so that in subsequent experimental cycles, the pipeline can better avoid proposing non-functional variants and instead focus the search on the most promising regions of the fitness landscape. These computational screening pipeline steps are shown in Figure 2.

**Figure 2.**
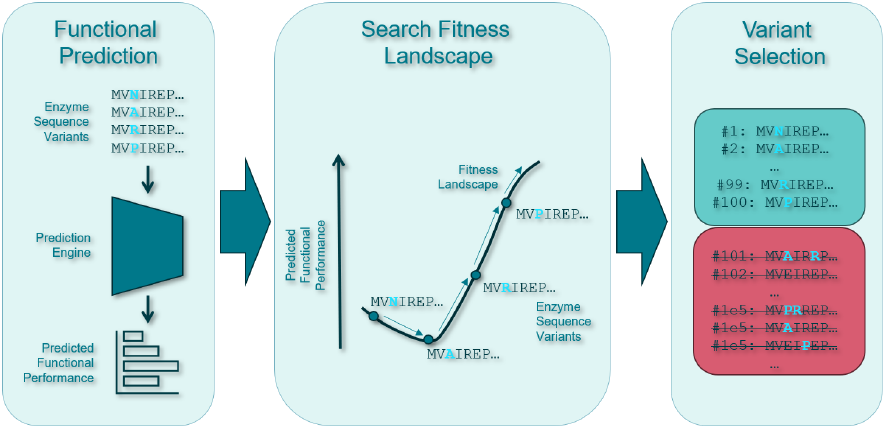
Details of the computational screening pipeline. The models predict the functional value for each variant sequence. The fitness landscape is searched for high value variants. At the final step, top variants are selected and passed to the laboratory team for expression and quantification.

### 1.2 Functional Performance

To tackle the challenge of predicting a protein variant’s functional properties, we utilized a family of advanced computational models called ESM (Evolutionary Scale Modeling) *protein language models* (PLMs) [1], [2]. These PLMs are based on transformer architectures, which were originally developed for natural language processing in *large language models* (LLMs) but have been adapted to understand the “language” of protein sequences.

The key advantage of the ESM models is that they can learn patterns and relationships in protein sequences without requiring any structural or functional data for training. Instead, they use a *self-supervised learning* approach, where the model is trained to predict deliberately masked amino acid residues within a large dataset of protein sequences. By learning to fill in these blanks, the model can capture complex patterns and long-range dependencies between amino acids that are far apart in the linear sequence but may be close together in the folded 3D structure of the protein.

This ability to learn intricate sequence patterns without relying on experimental functional data makes the ESM protein language models particularly well-suited for predicting the functional effects of mutations in proteins, even when little or no experimental data is available for a specific protein or variant.

The protein language models can be used to make predictions in two different ways, depending on the availability of functional data. The first approach is *zero-shot prediction*, which does not rely on any experimental functional data. Instead, the model makes predictions based solely on the patterns it learned during its self-supervised training on a vast number of naturally occurring protein sequences. For instance, if we change a single amino acid in a protein sequence, the model can predict whether the resulting mutant sequence is more or less likely to occur in nature, based on the evolutionary patterns it has learned. We can then use this likelihood as a rough proxy for the functional properties we’re interested in, assuming that these properties have been optimized by evolution.

The second approach, called *few-shot prediction*, becomes possible once we have a small amount of experimental functional data for a specific protein or set of variants. In this case, we use the pre-trained protein language model to convert each protein sequence into a numerical representation or *embedding*. This embedding captures the complex patterns and relationships within the sequence in a way that can be used as input for traditional machine learning models, such as linear regression or support vector machines. These models learn to predict the functional properties of interest from the embeddings, using the available experimental data as training examples. However, since the amount of experimental data is typically quite limited, we need to use specialized techniques like dimensionality reduction, regularization, and careful validation to ensure that the models can make accurate predictions across the entire fitness landscape, since the majority of sequences are not directly measured.

### 1.3 Fitness Landscape

Searching the fitness landscape for optimal protein variants presents several challenges. The first challenge is the sheer number of possible variants that can be generated from a given wild-type (WT) sequence. This number, denoted as *n*, can be calculated using the following combinatorial function:

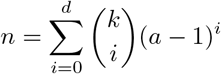

In this equation, *d* represents the number of positions in the sequence that are simultaneously mutated, *a* is the number of unique amino acids (usually 20), *k* is the length of the sequence, and 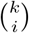 is the binomial coefficient. In the special case where *d* = *k*, then *n* = *a*^*k*^. Note this is assuming the only allowed variants are substitutions and excludes insertions or deletions.

Table 1 gives an example of how quickly the number of variants grows as a function of *d*. Note how between 1 and 2 mutations the number of variants increases 7, 500-fold, demonstrating how quickly the search space grows as we consider more simultaneous mutations. Exhaustively evaluating all possible variants becomes computationally infeasible for even moderate values of *d*. To overcome this challenge, we employ various search heuristics and optimization techniques that allow us to effectively sample the most promising regions of the fitness landscape without having to evaluate every possible variant.

**Table 1:** Number of possible protein variants for a given sequence length (*k*), number of amino acid alternates (*a*), and number of simultaneous mutations (*d*).

| Sequence Length<br>$k$ | Amino Acids<br>$a$ | Mutation Count<br>$d$ | Variant Count<br>$n$ |
| --- | --- | --- | --- |
| 800 | 20 | 1 | $1.52 \times 10^4$ |
| 800 | 20 | 2 | $1.15 \times 10^8$ |
| 800 | 20 | 3 | $5.83 \times 10^{11}$ |
| 800 | 20 | 4 | $2.21 \times 10^{15}$ |
| 800 | 20 | 5 | $6.68 \times 10^{18}$ |
| 800 | 20 | 6 | $1.68 \times 10^{22}$ |
| ... | ... | ... | ... |
| 800 | 20 | 800 | $20^{800} \approx 10^{1041}$ |

One way to search this vast fitness landscape is to start with an initial sequence and iteratively mutate it, generating a new variant at each step. If the new variant performs better than the previous one, we keep it and use it as the basis for the next round of mutation. However, if the new variant performs worse, we may decide to still keep it depending on an *acceptance probability*.

This probabilistic acceptance of worse variants allows the search process to escape from local optima, which are peaks in the fitness landscape that are higher than their immediate surroundings but lower than the global optimum. By occasionally accepting worse variants, the search can explore more of the rugged fitness landscape and potentially discover higher peaks elsewhere.

This type of heuristic search is known as a *Monte Carlo Markov chain* (MCMC), and it uses the *Metropolis-Hastings update rule* to determine the probability of accepting a new variant. The acceptance probability *p*_accept_ is defined as:

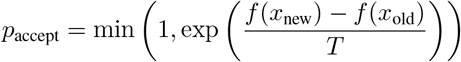

Where *f* (*x*_new_) and *f* (*x*_old_) are the functional value of the new and previous variants. The *T* parameter is a global “temperature” that controls the overall willingness to accept worse variants.

A technique called *simulated annealing* involves gradually decreasing this temperature parameter over time, making the search more likely to accept worse variants early on and then increasingly favoring better variants as the search progresses. This allows the search to initially explore a broad region of the fitness landscape and then gradually focus in on the most promising areas, ultimately settling into a near-optimal solution.

Another challenge in searching the fitness landscape arises from the uncertainty in our predictions. When we use protein language models to predict the performance of variants, we typically get a single predicted value, called a *point estimate*, for each variant. However, relying solely on these point estimates can result on focusing too much on a few promising areas and fail to explore other potentially valuable regions of the landscape. This is particularly problematic early on in a protein engineering campaign, when our predictions are likely to be less accurate. To address this issue, we need to find a balance between exploring new areas of the landscape, which may have high potential but also high uncertainty, and exploiting the best variants found so far, which have lower uncertainty but may not be optimal. This balance is known as the *exploration-exploitation trade-off*. One way to quantify this trade-off is through Bayesian optimization, a technique that incorporates uncertainty estimates into the search process.

In Bayesian optimization, instead of just predicting a single performance value for each variant, we also estimate the uncertainty around that prediction. This allows us to calculate an optimistic estimate of performance using the *upper confidence bound* (UCB) and a pessimistic estimate using the *lower confidence bound* (LCB). By favoring variants with high UCB values in the early experimental cycles of the campaign, we can prioritize exploration of promising but uncertain regions. At later interactions, when our predictions become more reliable, we can shift towards variants with high LCB values to exploit the best-performing variants with lower risk.

Furthermore, we can incorporate an additional prediction for each variant, the probability that it will be completely inactive. By multiplying this probability by the UCB, we get an “expected UCB” that adjusts the optimistic performance estimate based on the likelihood of the variant being non-functional. This helps us avoid wasting resources on variants that may have high predicted performance but are likely inactive.

To create a function that incorporates uncertainty, we use *Gaussian Processes* (GPs). A GP is a probabilistic model that fits a distribution of functions to the data, such that for a given input, the output is a Gaussian distribution instead of a single point estimate. This output distribution provides us with both a mean prediction and a measure of uncertainty, enabling us to use the Bayesian optimization techniques discussed above.

## 2 Literature Review

Protein engineering, a field that aims to create proteins with enhanced or novel properties, has traditionally relied on two approaches, namely *rational design* and *directed evolution* [3]. These engineered proteins find applications in various areas, such as medicine, industrial catalysis, and environmental remediation. Rational design is based on scientific understanding and involves modifying specific amino acids to improve properties like stability, activity, or specificity. Since this method depends on manually designing variants, the process is slow and error-prone. On the other hand, directed evolution involves random mutation of protein sequences, which is followed by characterization and selection of the best-performing variants. The best performing variants are then selected and used as a starting point for the next round of mutations. The process is iterated until the desired properties are obtained [4]. Despite its effectiveness, directed evolution is expensive and time-consuming, as it requires high-throughput processing of a large number of modified variants.

### 2.1 Models

Recent advancements in deep learning have revolutionized the field of protein engineering. A family of models called *protein language models* (PLMs) have emerged, adapting representation learning approaches from *natural language processing* (NLP) and *large language models* (LLMs) to learn from amino acid sequences and generate numeric representations. These models have shown remarkable success in various protein engineering tasks, such as predicting the impact of mutations on protein function and stability. This project evaluates current state-of-the-art representation learning models. An excellent systemic survey by Zhang et al. examines the lineages of LLM research in the biology and chemistry domains, including PLMs [5].

One of the earliest successful implementations of PLMs for protein variant selection was UniRep, an LSTM-based model. UniRep’s performance was validated by engineering and testing physical versions of the top-performing protein sequences in the lab [6]. A subsequent study demonstrated few-shot learning capabilities of UniRep, producing successful protein variants in wet lab validation [7]. Building upon this foundation, successive transformer-based models, such as TAPE, ESM-1b, ESM-1v, ProtTrans, and ESM-2, have achieved state-of-the-art results, showcasing improved performance in various protein engineering tasks [1], [2], [8], [9], [10]. Among these models, ESM-1v is specifically trained for engineering variants of a wild-type protein.

To further enhance the performance of PLMs, researchers have incorporated additional biological context. For instance, ECNet leverages *multi-sequence alignments* (MSA) of related proteins to provide the model with evolutionary context [11]. Similarly, LM-GVP employs graph neural networks to enable language models to learn structural information of proteins [12].

A recent breakthrough in PLMs is ESMFold, which can predict the three-dimensional structure of a protein using only its amino acid sequence [2]. This is a significant advancement compared to AlphaFold2 [13], which requires additional information such as MSA, residue pair representation, and template structures. ESMFold achieves similar or better performance than AlphaFold2, highlighting the potential of PLMs in protein structure prediction [2]. Accurate structure prediction is crucial for understanding protein function and designing novel proteins with desired properties.

In addition to protein variant selection and structure prediction, PLMs have also been applied to generating novel proteins. Generative models, such as *Generative Adversarial Networks* (GANs) and *Variational Auto-Encoders* (VAEs), have shown success in this domain. For example, a GAN-based approach has been used to generate novel functional proteins [14], while DeepSequence, a VAE-based model, has demonstrated the ability to generate novel proteins with desired properties [15].

Interestingly, the ESMFold PLM, originally designed for structure prediction, has also been adapted for protein generation. It can perform both unconstrained and *fixed backbone generation* [16]. In fixed backbone generation, the backbone atom positions are predetermined, and the model learns which amino acid side chains will result in the given conformation. ESMFold can also incorporate physical constraints, such as hydrophobicity, into the optimization process [17].

A promising new class of generative models called *diffusion models* has recently emerged. These models are trained by taking a known protein structure, incrementally adding noise, and then learning to *denoise* the structure. This round-trip process allows diffusion models to achieve state-of-the-art performance in generating novel protein structures from pure noise [18]. This approach opens up new possibilities for designing proteins with desired structural and functional properties.

Once a model has been selected to predict the functional performance of protein variants, it can be used to explore the *fitness landscape* and identify superior variants. The fitness landscape represents the functional relationship between a protein’s sequence and its corresponding functional value, such as catalytic activity or stability. Navigating this landscape efficiently is crucial for discovering optimal protein sequences.

Several optimization methods have been employed to explore the fitness landscape. Monte Carlo Markov Chain (MCMC) methods have been used to sample the landscape and identify high-performing variants [7]. Bayesian optimization, another popular approach, balances exploration and exploitation to efficiently search the landscape and find optimal sequences [19], [20], [21]. This project investigates both MCMC and Bayesian optimization techniques to compare their effectiveness in protein engineering tasks.

In addition to directly optimizing the protein sequence, optimization can also be performed in the latent space of embeddings learned by the model. For example, a transformer-based auto-encoder model called ReLSO has been used to optimize protein sequences in the embedding space [22]. This approach allows for a more compact representation of the sequence space and can potentially lead to more efficient optimization.

### 2.2 Data

Training PLMs using self-supervised learning requires large and diverse datasets. Two commonly used datasets for this purpose are UniRef50 and UniRef90 [23]. These datasets are constructed by clustering protein sequences based on their sequence identity, with UniRef50 and UniRef90 containing clusters of sequences with more than 50% and 90% sequence identity, respectively. Each cluster is represented by a single sequence in the dataset, which helps maintain balance and reduce redundancy. Currently, UniRef50 contains 59 million sequences, while UniRef90 contains 166 million sequences. The choice of dataset can impact model performance on different tasks. Models trained on UniRef90 typically perform better on variant selection tasks, while those trained on UniRef50 excel in structural prediction tasks. In addition to sequence data, structural models such as AlphaFold2 and ESMFold also utilize the Protein Data Bank (PDB) [24], a repository of over 200,000 experimentally-determined protein structures obtained through methods like X-ray crystallography.

While sequence data is abundant, sequence-to-function data, which maps protein sequences to their functional properties (e.g., enzymatic activity or thermal stability), is scarce for most proteins. However, for certain well-studied model proteins, *deep mutational scanning* (DMS) datasets are available in the literature. DMS studies systematically mutate and characterize a large number of protein variants to assess the impact of mutations on protein function, comparing them to the wild-type (WT) sequence. These studies also investigate the effects of *epistasis*, which refers to the non-linear interactions between multiple mutations and their influence on functional performance [15], [25], [26], [27]. The Data Sources section of this report provides more details on the specific datasets used in this project.

## 3 Methods

### 3.1 Data Sources

The dataset used in this study consists of 34 deep mutational scanning (DMS) studies[15], [25], [26], [27] that quantify the impact of mutations on protein function compared to their respective wild-type (WT) sequences. These studies span 25 organisms, including viruses, bacteria, jellyfish, and humans. The number of variants per study ranges from 300 to 12,000, with protein sequence lengths varying from 72 to 881 residues. To facilitate comparison across studies, the functional data has been log-normalized relative to the WT value for each protein. Supplemental Table S1 provides further details on the individual DMS datasets used in this analysis.

### 3.2 Zero-shot Learning

In this study, we employ the Evolutionary Scale Modeling (ESM) family of transformer-based PLMs developed by the Facebook AI Research group (FAIR). FAIR^*^ has released ESM-1 [9] and ESM-2 models [2], along with several task-specific variants. Table S2 provides a list of the various ESM models available, and the code for these models can be found at https://github.com/facebookresearch/esm.

The initial paper introducing the ESM-1v models [1] demonstrated their effectiveness in zero-shot learning. ESM-1v is an ensemble of five transformer-based PLMs, each containing 650 million parameters (see Supplemental Table S2 for more details). To make zero-shot predictions, the models compare the likelihood of the variant sequence to that of the WT sequence. For variants with multiple mutations, a marginal method is required to aggregate the likelihoods. In our project, we implemented the wild-type marginal due to its fast inference speed and performance comparable to the mutation marginal. The original paper provides more details on the marginals and their trade-offs.

We performed zero-shot inference on all 34 protein datasets, limiting each to 1,000 variants. These variants were then passed through each of the five ESM-1v models, and the predicted WT marginal likelihoods were averaged. To assess the model’s performance, we calculated Spearman’s rank correlation between the averaged likelihoods and the functional values. Spearman’s correlation is used because our objective is to predict the top-performing variants, regardless of the magnitude of the predicted values.

### 3.3 Restricted Fitness Landscape Search

To evaluate the effectiveness of the zero-shot predictions in identifying superior variants, we performed a restricted fitness landscape search. Since only 1,000 variants were predicted for each protein, we treated this as a simulated search of a known landscape. The goal is to demonstrate how quickly superior variants can be discovered using the zero-shot predictions compared to random selection.

For each of the 34 proteins, we conducted a simulation comparing a zero-shot prediction strategy to a random selection strategy. In each round of the simulation, a variant is selected, and its true functional value is checked. The simulation records the number of rounds needed to find half of the top 10% of variants based on their functional values, which we refer to as the *rounds to half of the top decile* (RHTD). The zero-shot strategy always selects the variants with the top 1% of predicted values in each round, while the random strategy randomly selects 1% of variants without replacement.

To estimate the distribution of RHTD, we performed 1,000 independent runs for each protein, with each run limited to an independent sub-sample of 100 variants from the available 1,000. This approach allows us to assess the robustness of the zero-shot predictions in identifying superior variants across different subsets of the fitness landscape. A RHTD ratio is calculated by dividing the PLM RHTD by the random RHTD. This ratio is the expected reduction in rounds using the zero-shot predictor.

### 3.4 Few-shot Learning

To evaluate few-shot learning, we selected several ESM PLMs to provide embeddings for regression models, which we refer to as *regressors*. We chose the first ESM-1v PLM with 650 million parameters and four ESM-2 PLMs with 8, 35, 150, and 650 million parameters each. The 3 and 15 billion parameter PLMs were not included due to GPU RAM constraints. These PLMs were used to generate numeric embeddings of the sequences for each protein dataset, with embedding dimensions ranging from 320 to 1280. The embeddings serve as the feature set for the regressors. Supplemental Table S2 provides more details on the embedding models. To avoid excessive computation, embeddings were often limited to 1,000 per dataset.

Four regressors were selected to predict the functional values of each protein dataset using the numeric embeddings: *K-Nearest Neighbor, Support Vector Machine, Random Forest*, and *Ridge Regression*. Each regressor employs a different mechanism to fit the data, allowing for a diverse examination of techniques. Grid search with 5-fold cross-validation is used for hyperparameter tuning to select optimal hyperparameters while minimizing overfitting. Spearman rank correlation is used as the evaluation metric during hyperparameter tuning, and the best-performing model is selected for further evaluation.

To assess regressor performance with limited data, we trained the regressors with 30, 100, and 300 data points. In all these scenarios, the PLM embedding sizes exceed the number of training data points. To improve the fit, *Principal Component Analysis* (PCA) is employed to reduce the dimensionality of the embeddings. The number of principal components examined is 4, 10, and 24. To account for training variance, three replicates of each training instance were performed using different seeds for data splitting and initializing the regressor states.

In total, 4,590 training combinations were examined for a total of 18,360 trained regressors. For each of these instances, the best-performing model from hyperparameter tuning is evaluated by calculating the Spearman rank correlation on the holdout data.

### 3.5 Few-Shot Gaussian Processes

The fitting of Gaussian Process (GP) regressors to the datasets followed a similar approach to that used for the few-shot regressors. Hyperparameter tuning is employed to select the best-performing kernels. The same PLM embeddings, protein datasets, training sizes, principal components, and replicate seeds is also used. Spearman rank correlation is used for both hyperparameter tuning and holdout data evaluation.

In addition to GP regressors, a set of GP classifiers is fit for each training experimental cycle. These classifiers predict whether a given variant will exceed a threshold, which is particularly useful for bimodal datasets containing both high and low functional values, such as variants that fail to express or lose enzymatic activity. To account for the diversity of the datasets and potential censoring of low functional values, we developed an automated method to identify a bimodal threshold. This method involves fitting a *kernel density estimate* (KDE) to the functional values and identifying the KDE peaks using a peak finding algorithm from signal processing. The threshold is set to the minimum value between the first two peaks. If no peaks are found, the threshold is set to the WT functional value. In cases where either of these thresholds results in an imbalance of more than 70% of the values, the threshold is set to the 50th percentile.

The GP regressors return both a mean (*µ*_*GP*_) and a standard deviation (*σ*_*GP*_) value for each data point, while the GP classifiers return the probability (*p*_*GP*_) of a given data point belonging to the high-functioning mode. Using these outputs, we calculate four predictors (*ŷ*):

The mean value of the GP regressor (*ŷ*_*P E*_), analogous to a point estimate from conventional regressors:

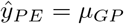

The Upper Confidence Bound (UCB) (*ŷ*_*UCB*_), which accounts for the potential of superior variants due to high estimates and high uncertainty:

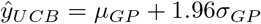

The expected UCB (*ŷ*_*E*(*UCB*)_), which accounts for the probability of the variant being high-functional:

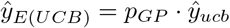

For comparison, the ridge regressor model described earlier is recalculated alongside the GPs to serve as a baseline.

### 3.6 Bayesian Optimization

A simplified Bayesian optimization procedure is performed by using the GP predictors on the restricted fitness landscape search method used for the zero-shot prediction task. This simplified approach assumes 300 training data points and then fits the GP regressors per the few-shot GP procedure. The same series of simulations are then performed to calculate the RHTD values for the expected UCB, UCB, point estimate, and random cases. RHTD ratios are calculated for each of the three GP predictors versus the random baseline case.

A simplified Bayesian optimization procedure is implemented by applying the Gaussian process (GP) predictors to the restricted fitness landscape search method used for the zero-shot prediction task. This approach assumes the availability of 300 training data points and fits the GP regressors according to the few-shot GP procedure described earlier. A series of simulations is then performed to calculate the RHTD values for the expected UCB, UCB, point estimate, and random selection cases. RHTD ratios are computed for each of the three GP predictors by dividing their respective RHTD values by the RHTD value of the random baseline case. These ratios provide a measure of the improvement in search efficiency achieved by the Bayesian optimization approach compared to random selection.

### 3.7 Software and Computation

The analysis is conducted using Python, with the ESM models implemented in the PyTorch framework. Other Python libraries used in this study include NumPy, pandas, SciPy, and scikit-learn. Inference with the ESM models was performed on a GPU, while training and inference of the scikit-learn supervised models were executed on a CPU. The analysis code and package requirements are available upon request.

Computations were performed on a Dell Precision 7680 laptop equipped with an Intel Core i7-13850HX processor (20 cores, 2.1 GHz), 64 GB of RAM, and an NVIDIA RTX 2000 GPU with 8 GB of dedicated RAM.

When working with ESM models, it is crucial to consider the batch size used for a given model. The GPU memory allocates space for the model weight parameters and the tokens for each batch. As the model size increases, the batch size needs to be decreased accordingly. Exceeding the available GPU memory can lead to a severe decrease in processing speed and should be avoided. In this study, the ESM-2 model variants with 3B and 15B parameters were excluded due to the inability to load them without exceeding the available GPU memory. Performing inference on a GPU provides a 5x to 20x speedup compared to running the same tasks on a CPU.

## 4 Results

### 4.1 Zero-shot Learning

The zero-shot predictor demonstrated a strong correlation between the top predicted variants and the top experimentally validated variants. Figure 3 illustrates this correlation for the `AMIE PSAEA Whitehead` protein dataset, revealing a clear positive trend between the experimental rank and the predicted rank. Figure 4 shows that 31 out of the 34 protein datasets exhibit high Spearman rank correlations (*ρ* > 0.2), with a median value of *ρ* = 0.44 and a mean value of *ρ* = 0.50. These results are particularly impressive considering that the predictions are made without any functional training data. Two of the proteins had *ρ* ≈ 0, indicating no significant correlation between the predicted and experimental ranks. Interestingly, one protein had a *ρ* ≈ − 0.5, suggesting that the functional value being selected may violate the assumption that higher functional values align with evolutionarily optimal mutations. This finding warrants further investigation to understand the underlying factors contributing to this discrepancy.

**Figure 3.**
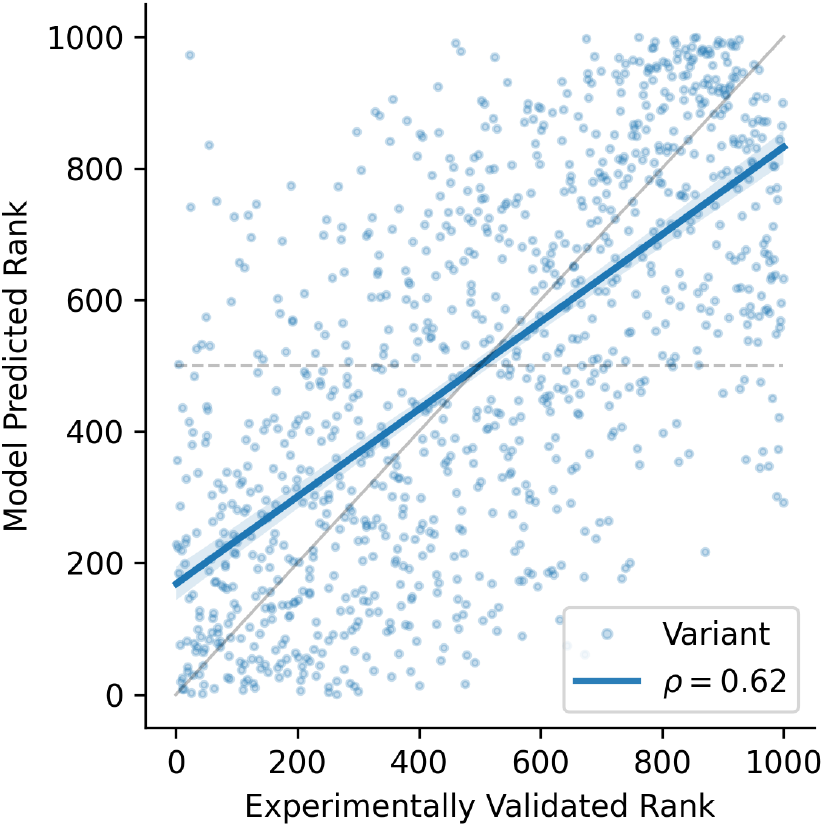
Visualization of Spearman correlation (*ρ*) for the zero-shot prediction. Each variant is plotted according to the rank of its experimentally validated functional value versus the rank of the predicted functional value. The calculated correlation is proportional to the slope of the blue fitted line. The solid grey line and dashed gray line correspond to *ρ* = 1 and *ρ* = 0, respectively. This example is from `AMIE PSEAE Whitehead`.

**Figure 4.**
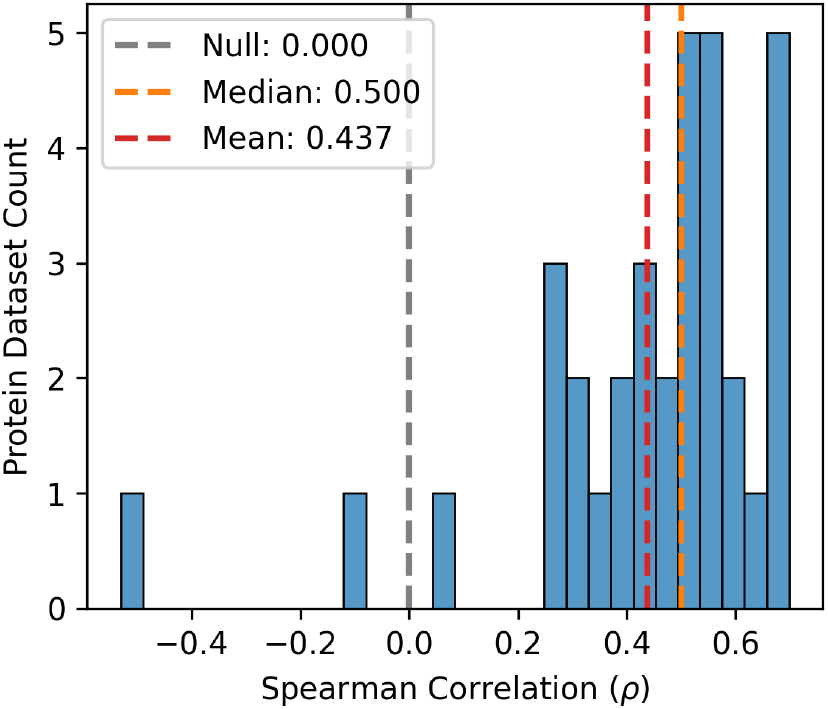
Distribution of the zero-shot prediction correlations for all 34 protein datasets.

### 4.2 Restricted Fitness Landscape Search

To investigate the impact of high correlations on fitness landscape search, a series of simulations were performed. The *rounds to half of the top decile* (RHTD) metric quantifies the number of variants that must be selected to obtain half of the top 10% of the population. Figure 5 illustrates an example from the `AMIE PSEAE Whitehead` protein dataset. The random selection strategy requires an average of 47 rounds, while the zero-shot predictions from the ESM1v PLM achieve the same goal in only 25 rounds on average.

**Figure 5.**
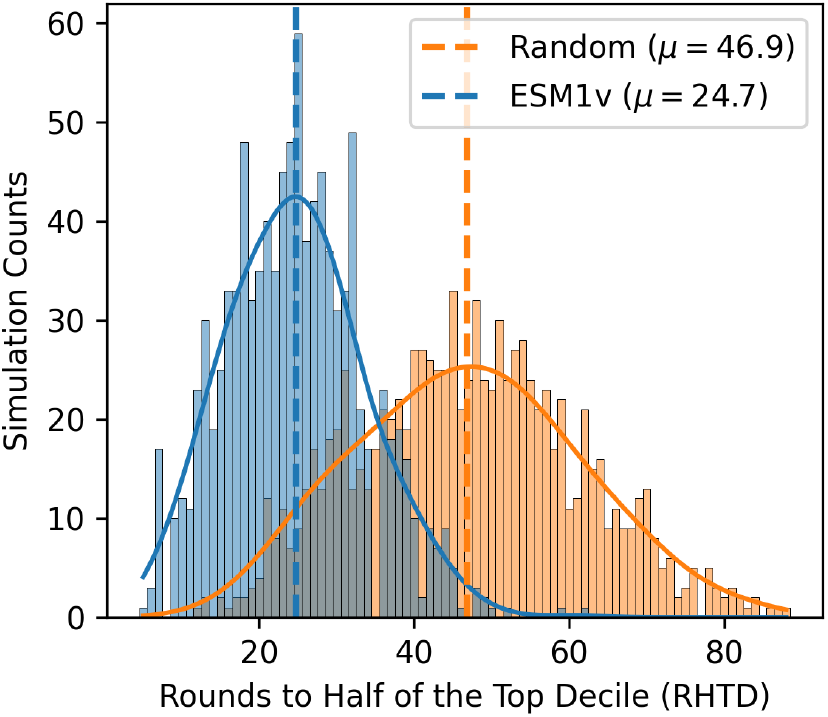
The distribution of the *rounds to half of the top decile* (RHTD). This metric represents the number of rounds needed to select half of the top 10% of variants. The ESM1v model values are shown in blue, while the random selection values are displayed in orange. This example is based on the `AMIE_PSEAE_Whitehead` protein dataset with 1,000 independent simulations.

The RHTD ratio is employed to measure the reduction in the number of rounds compared to random variant selection. Figure 6 presents the RHTD ratios for each simulation run across the protein datasets. The mean RHTD ratio is 0.69, and the median RHTD ratio is 0.55, suggesting that a typical reduction of more than 45% can be expected. However, there is some risk involved, as 18% of the simulations yield RHTD ratios greater than 1, indicating worse performance than random selection.

**Figure 6.**
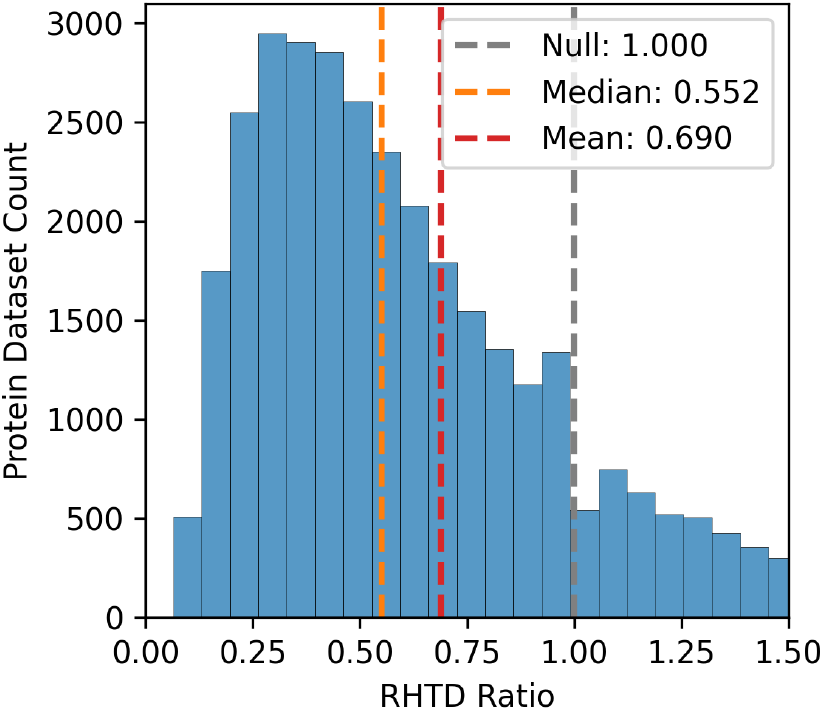
The distribution of the RHTD Ratio, which is calculated by dividing the RHTD of the ESM1v model by the RHTD of random selection. The null case, where the RHTD Ratio equals 1, is shown in grey. The mean and median RHTD Ratios are indicated in orange and red, respectively.

Figure 7 explores the relationship between Spearman correlation and the RHTD ratio. Each data point represents the mean correlation and RHTD ratio values for a specific protein dataset. The figure reveals a strong, albeit imperfect, correlation between these two measures. This finding suggests that while the Spearman correlation is a good indicator of the reduction in selection effort for a given model and dataset, it is not a perfect predictor.

**Figure 7.**
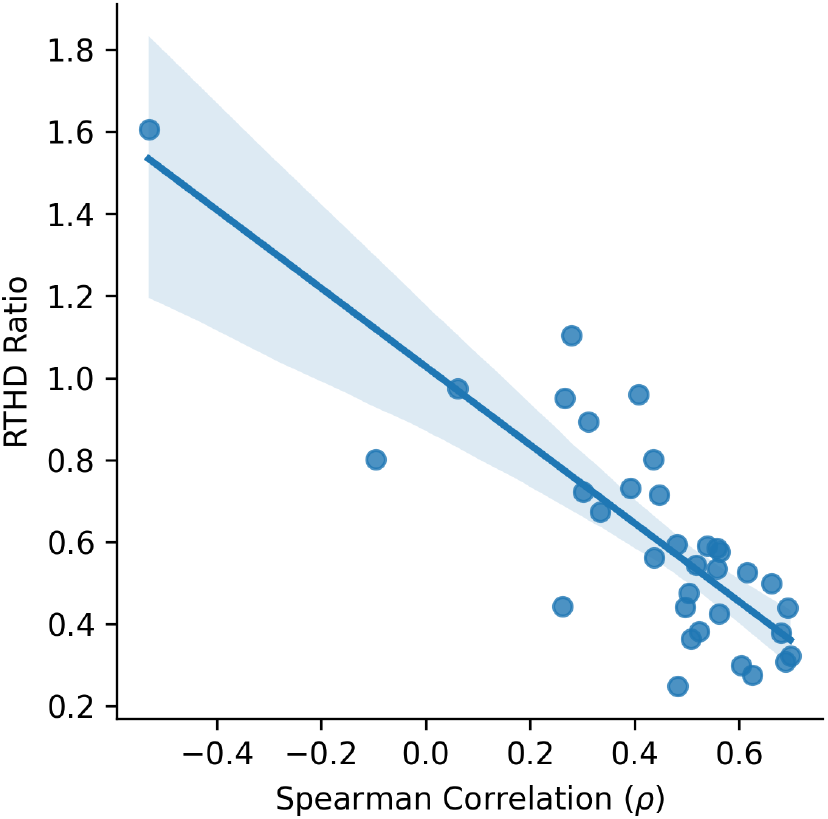
A scatter plot illustrating the relationship between the zero-shot Spearman correlation and the mean RHTD Ratio. Each data point corresponds to a protein dataset. The trend line is fitted to the data points and includes a 95% confidence interval estimate.

### 4.3 Few-shot Learning

In addition to the zero-shot predictions, the performance of few-shot predictions was analyzed. The predictions were made by training regression models on the embeddings obtained from the PLMs. The types of regressors used include k-nearest neighbors, support vector machine (SVM), random forest, and ridge regression. These models were trained on datasets of 30, 100, and 300 data points, using 4, 10, and 24 principal components as the feature set. Figure 8 shows the performance of each regressor type for the ESM1v PLM with a 24 principal component feature set. Both the k-nearest neighbors and the random forest regressors underperform across all training sizes. The ridge regressor outperforms other models at smaller training sizes, while the SVM regressor exhibits superior performance at larger training sizes. This may indicate that the ridge regressor better generalizes with few data points due to its regularization technique, whereas SVM achieves better generalization when more data is available.

**Figure 8.**
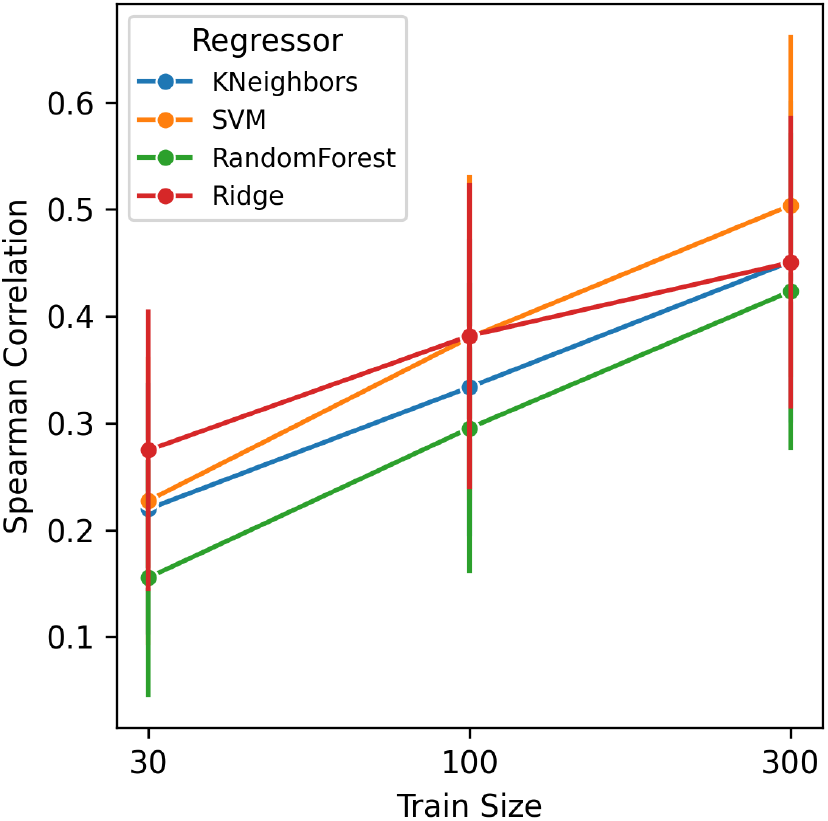
A line plot with error bars for few-shot regression results. The colors denote the regressor models across the three training data sizes. The y-axis represents the Spearman correlation on the holdout data. The results are filtered for the ESM1v PLM with 24 principal components.

A comparison of the PLMs that provide embeddings to the regressors is shown in Figure 9. The model parameter size is monotonically related to performance, with saturation being reached by the PLMs with ≥ 150M parameters. The ESM1v 650M and ESM2 650M models exhibit nearly identical performance. The ESM1v was trained on the UniRef90 data, which reportedly provides more nuanced information for variant prediction, while the ESM2 was trained on the UniRef50 data, which performs better on other tasks. ESM2 has an improved architecture that yields higher performance compared to ESM1 UR50 models. These trade-offs seem to balance out in this analysis.

**Figure 9.**
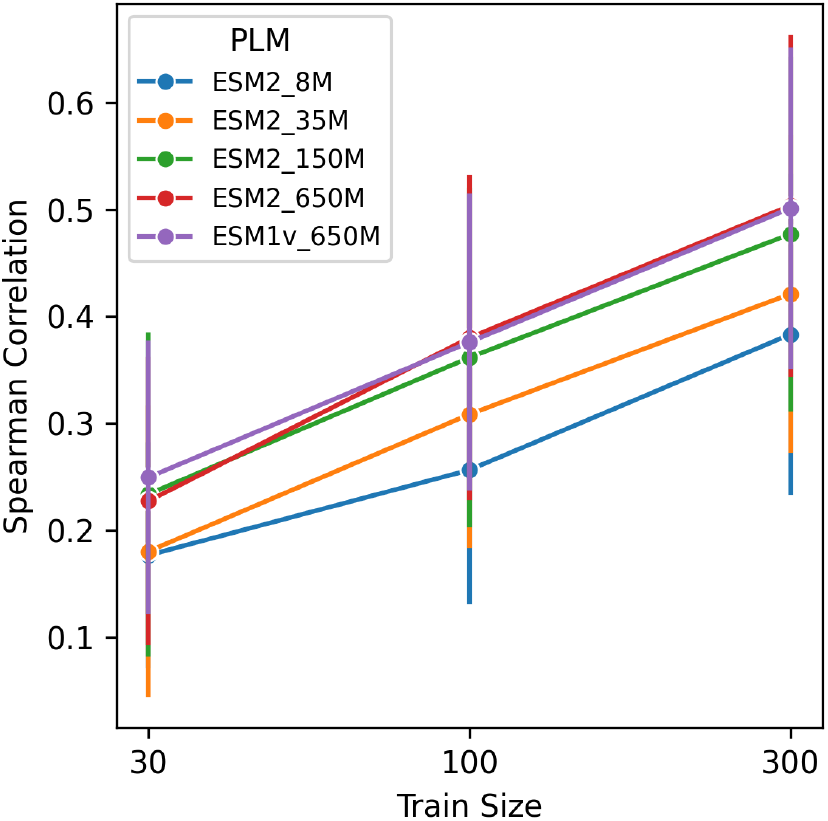
A line plot with error bars for few-shot regression results. The colors denote the PLM models across the three training data sizes. The y-axis represents the Spearman correlation on the holdout data. The results are filtered for the SVM regressor with 24 principal components.

A more detailed breakdown of the Spearman correlations for each training size is shown in Figure 10. This case corresponds to SVM regressors using 24 principal components, with results of the three replicates across the 34 protein datasets. The mode of each training size increases, with *ρ* ≈ 0.50 for 300 training data points. Even with 300 data points, a small minority of results show low correlation, indicating some risk of protein functional values with low predictive power.

**Figure 10.**
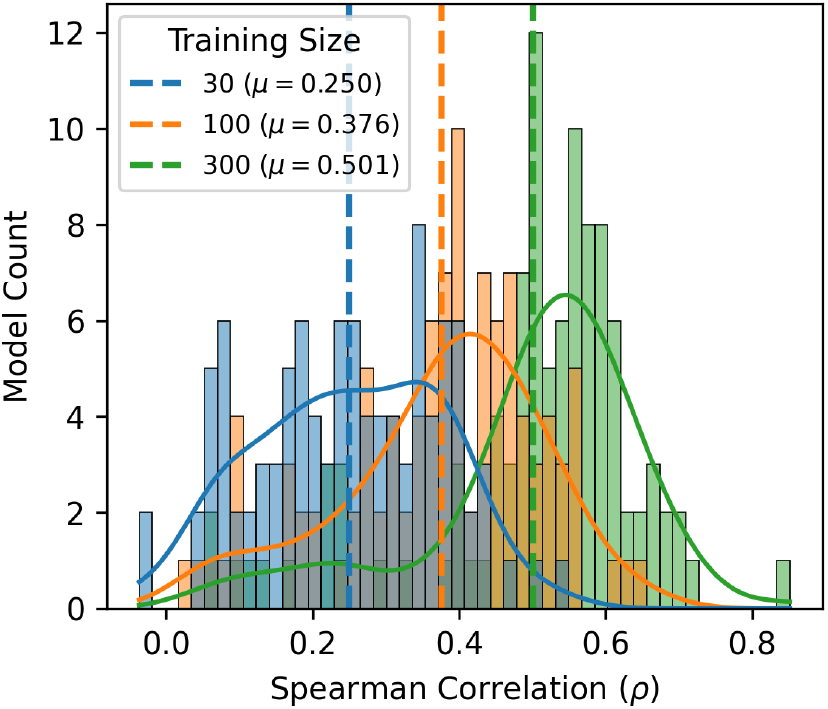
A series of histograms for few-shot regression results, showing distributions of the ESM1v PLM and the SVM regressor selected for 24 principal components. The colors correspond to the three different training data sizes. The x-axis represents the Spearman correlation on the holdout data. The variation is across the 34 protein datasets and 3 replicates.

### 4.4 Gaussian Processes

In addition to regressors that provide point estimates, we also explored GPs to estimate a normal distribution for each variant prediction. We trained a GP classifier to predict the probability of highly functional variants. From the GP regressor and classifier, we constructed three predictors: the point estimate, the UCB, and the expected UCB. Figure 11 shows the Spearman correlation for each predictor, as well as for a ridge regressor from the previous section, which is included as a baseline. The ridge regressor performs best at the small training size, while the GP predictors outperform at larger sizes. The point estimate and expected UCB even surpass the performance of the SVM regressor from the previous section.

**Figure 11.**
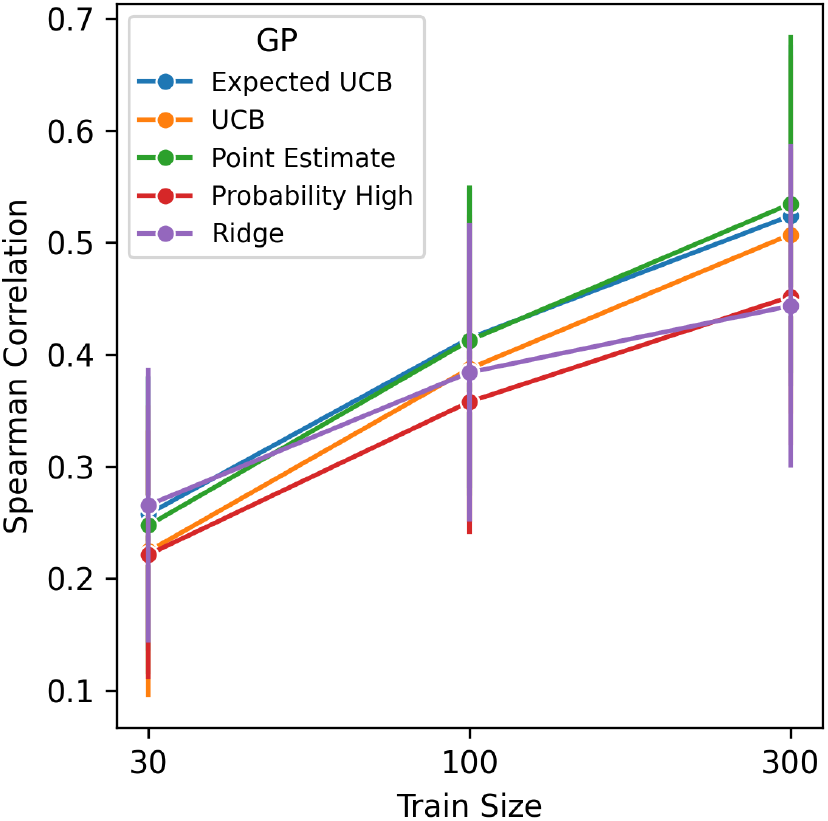
A line plot with error bars for few-shot Gaussian process predictors. The colors denote the GP predictors across the three training data sizes. The y-axis represents the Spearman correlation on the holdout data. The results are filtered for the ESM1v PLM with 24 principal components.

The PLM models are also compared for the GP predictors, as shown in Figure 12. Similar to the previous regressors, ESM2 650M and ESM1v 650M exhibit nearly identical performance, while other models underperform. A breakdown of the Spearman correlations for each training size is shown in Figure 13. This case corresponds to expected UCB predictors using 24 principal components, with results of the three replicates across the 34 protein datasets. The mode of each training size increases, with *ρ* ≈ 0.52 for 300 training data points, which is a small improvement over the SVM regressor in Figure 10.

**Figure 12.**
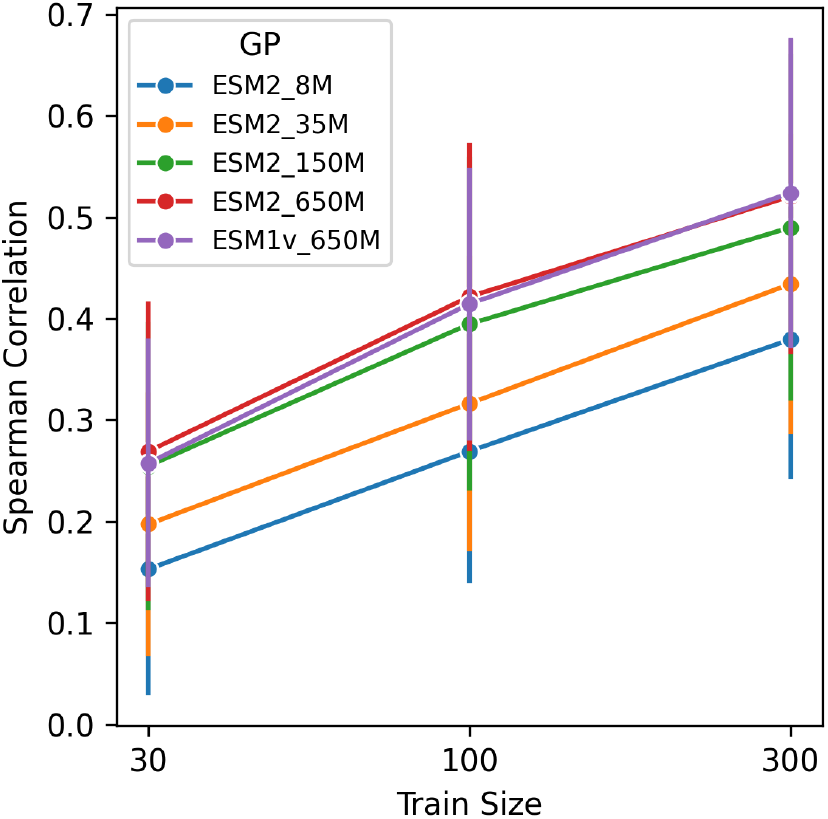
A line plot with error bars for few-shot Gaussian process predictors. The colors denote the PLM models across the three training data sizes. The y-axis represents the Spearman correlation on the holdout data. The results are filtered for the Expected UCB predictor with 24 principal components.

**Figure 13.**
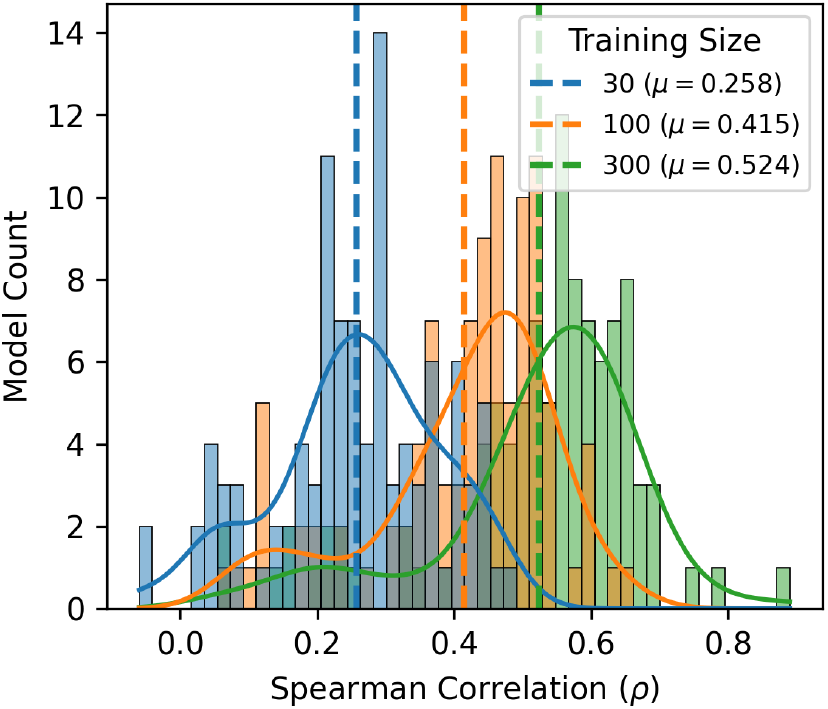
A series of histograms for few-shot Gaussian process predictors, showing distributions of the ESM1v PLM and the Expected UCB predictor selected for 24 principal components. The colors correspond to the three different training data sizes. The x-axis represents the Spearman correlation on the holdout data. The variation is across the 34 protein datasets and 3 replicates.

### 4.5 Bayesian Optimization

Similar to the fitness landscape search for the zero-shot predictions, the GP predictors are used to leverage their uncertainties in searching the landscape. This approach is known as Bayesian optimization (BO). Our implementation is simplified and does not retrain the GP at each selection round, providing a consistent comparison to the zero-shot simulations. The BO simulations use the training data with 300 points and 24 principal components.

Figure 14 shows the performance of the expected UCB and the point estimate predictors, which have mean RHTDs of 25.5 and 22.9, respectively. The point estimate outperforms the expected UCB, which may indicate that the GP requires more data to better estimate the uncertainties. The point estimate also outperforms the zero-shot case from Figure 5, which has a mean of 24.7.

**Figure 14.**
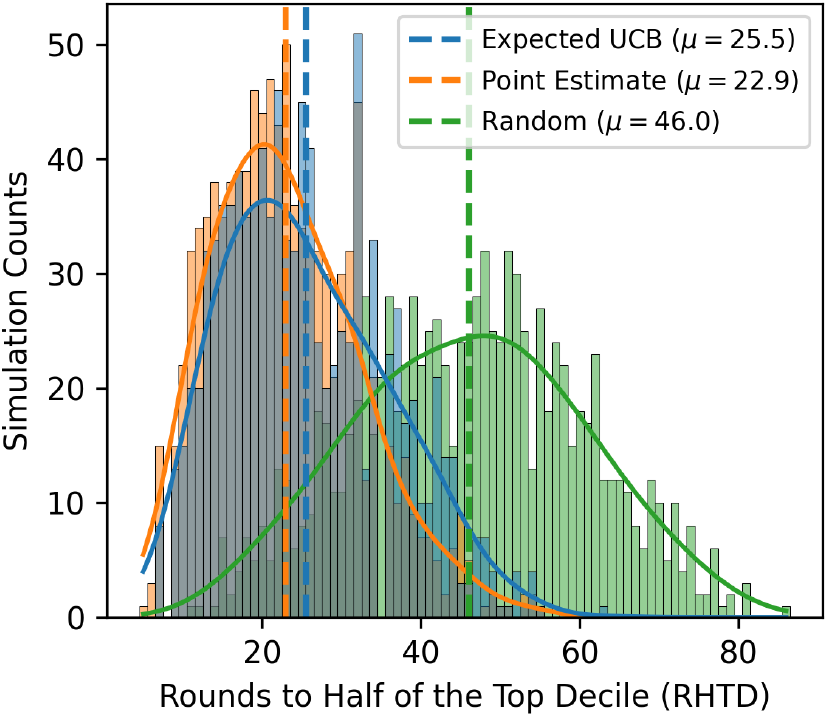
The distribution of the *rounds to half of the top decile* (RHTD) for Bayesian Optimization. This represents the number of rounds needed to select half of the top 10% of variants. The color code corresponds to the GP predictors, with random selection values shown in orange. This example is based on the `AMIE PSEAE Whitehead` dataset.

The calculated RHTD ratios distribution is shown in Figure 15. The mean and median values are 0.58 and 0.48, respectively, compared to 0.69 and 0.55 for the zero-shot case in Figure 6. This represents a substantial improvement and further reduces the cost and effort of protein engineering campaigns. The risk of having an RHTD ratio ¿1 is also reduced from 18% to 10%.

**Figure 15.**
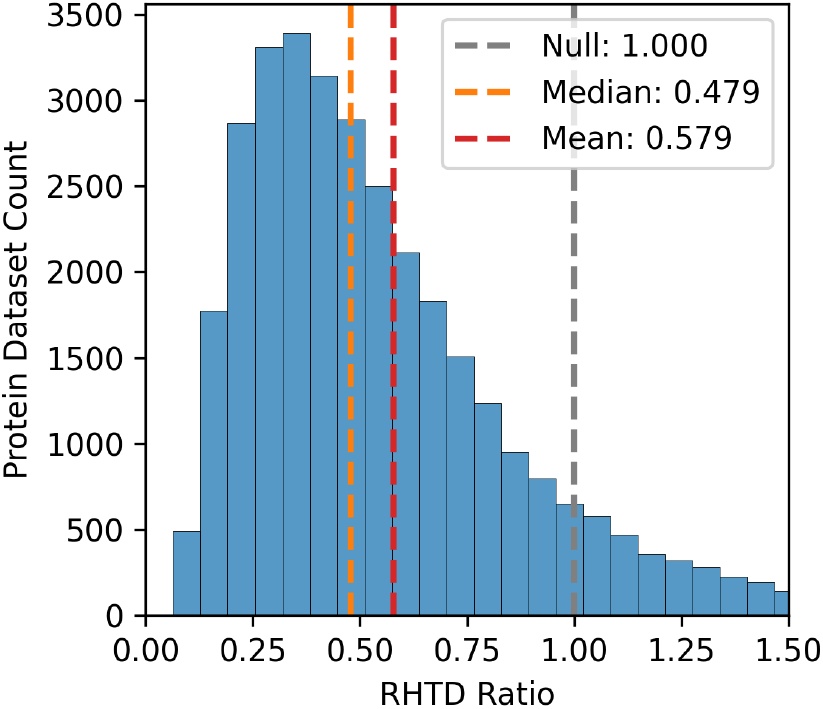
The distribution of the GP point estimate predictor RHTD Ratios, calculated as the GP RHTD divided by the random RHTD. The null case, where the RHTD Ratio equals 1, is shown in grey. The mean and median RHTD Ratios are indicated in orange and red, respectively.

## 5 Discussion

The use of both zero-shot predictions from PLMs and few-shot predictions with GPs shows great promise in reducing the cost and effort associated with developing improved protein variants. When no data is available, zeroshot prediction is capable of reducing the required effort by nearly half. As more functional data becomes available, GPs can be employed to generate more accurate predictions and further enhance the ability to identify superior variants.

The GP predictions started to outperform the zero-shot predictions at 300 training data points. When data points are below 100, the zero-shot is superior. In practice, between 100 and 300 data points it may be best to phase out the zero-shot by doing a weighted average of the two approaches, where both model weights sum to 1 and the GP weight is proportional to the distance between 100 and 300.

While the models explored had high predictive power for the majority of the proteins, a few proteins had low performance. These cases should be examined further to determine whether they are inherently challenging to predict or if the datasets are poorly aligned with the variant screening objective under study. This investigation could provide valuable insights into the limitations of the current approach and guide future improvements. Combining both zero-shot and GP predictors could potentially lead to even better predictions. A stacked model ensemble with a top-level model that selects how to blend the zero-shot and GP predictions based on the number of training points and other features could help address the challenge of determining when to use each approach. This ensemble method could leverage the strengths of both techniques and adapt to the available data, resulting in more robust and accurate predictions across a wider range of scenarios.

The Bayesian optimization performance of the uncertainty-based acquisition functions (UCB and Expected UCB) unexpectedly underperformed relative to the GP point estimates, with mean RHTD values of 25.5 for Expected UCB compared to 22.9 for point estimates. Several interconnected factors likely contribute to this counterintuitive result.

*Data limitations and uncertainty calibration* represent the most probable explanation. Gaussian Process uncertainty estimation requires substantial training data for proper calibration, particularly in early screening rounds when functional measurements are scarce. The high-dimensional nature of protein embedding spaces (E SM embeddings span 320-1280 dimensions) further exacerbates this challenge, as GP uncertainty estimation becomes increasingly unreliable in high-dimensional regimes due to the curse of dimensionality.

*Methodological considerations* may also play a role. The exploration-exploitation balance in UCB is governed by the *β* parameter, and our default parameterization may have been suboptimal for protein fitness landscapes. Additionally, standard GP assumptions of Gaussian-distributed residuals may be violated if protein fitness landscapes exhibit multi-modal or heavy-tailed distributions, leading to poorly calibrated uncertainty estimates.

*Search space characteristics* provide another important consideration. Unlike previous Bayesian optimization studies in protein engineering that focused on combinations of sequence fragments—resulting in substantially smaller and more structured search spaces—our approach targets single-residue mutations across entire protein sequences. This difference in search space size and complexity (20^*L*^ versus more constrained combinatorial spaces) may fundamentally alter the effectiveness of uncertainty-based exploration strategies, suggesting that protein engineering applications may require specialized acquisition functions tailored to these unique landscape characteristics.

## Author Contributions

J.M. conceived and designed the study, developed the computational pipeline, performed all analyses, and wrote the manuscript. P.S. provided methodological guidance and contributed to manuscript review. Both authors approved the final manuscript.

## Data and Code Availability

The protein datasets used in this study are publicly available from deep mutational scanning studies as cited. Due to the computational nature of this work conducted as part of academic coursework, the analysis code is not currently available in a public repository.

## Acknowledgments

J.M. is currently affiliated with Mayo Clinic, Rochester, MN. We thank the Georgia Institute of Technology School of Industrial and Systems Engineering for academic support during this research.

## 6 Supplemental Material

### 6.1 Datasets

The various datasets used are listed in Table S2.

### 6.2 ESM Models

The various models in the ESM family are listed in Table S2.

**Table S1:** Selected protein dataset.

| Dataset Name | Protein Length | Total Variants |
| --- | --- | --- |
| AMIE_PSEAE_Whitehead | 346 | 4477 |
| B3VI55_LIPSTSTABLE | 447 | 6530 |
| B3VI55_LIPST_Whitehead2015 | 439 | 6315 |
| BF520_env_Bloom2018 | 852 | 12236 |
| BG505_env_Bloom2018 | 860 | 12217 |
| BG_STRSQ_hmmerbit | 479 | 2634 |
| BLAT_ECOLX_Ostermeier2014 | 286 | 4594 |
| BLAT_ECOLX_Palzkill2012 | 286 | 4788 |
| BLAT_ECOLX_Ranganathan2015 | 286 | 4788 |
| BLAT_ECOLX_Tenaillon2013 | 286 | 949 |
| CALM1_HUMAN_Roth2017 | 149 | 1729 |
| DLG4_RAT_Ranganathan2012 | 724 | 1557 |
| GAL4_YEAST_Shendure2015 | 881 | 1104 |
| HG_FLU_Bloom2016 | 565 | 10336 |
| HSP82_YEAST_Bolon2016 | 709 | 4084 |
| IF1_ECOLI_Kishony | 72 | 1311 |
| KKA2_KLEPN_Mikkelsen2014 | 264 | 4240 |
| MK01_HUMAN_Johannessen | 360 | 4382 |
| MTH3_HAEAEESTABILIZED_Tawfik2015 | 330 | 1719 |
| P84126_THETH_b0 | 254 | 1519 |
| PABP_YEAST_Fields2013-singles | 577 | 1187 |
| PA_FLU_Sun2015 | 716 | 1820 |
| PTEN_HUMAN_Fowler2018 | 403 | 3012 |
| RASH_HUMAN_Kuriyan | 189 | 3078 |
| RL401_YEAST_Bolon2013 | 128 | 1159 |
| RL401_YEAST_Bolon2014 | 128 | 1293 |
| RL401_YEAST_Fraser2016 | 128 | 1199 |
| SUMO1_HUMAN_Roth2017 | 101 | 1328 |
| TIM_SULSO_b0 | 248 | 1519 |
| TIM_THEMA_b0 | 252 | 1519 |
| TPK1_HUMAN_Roth2017 | 243 | 2591 |
| TPMT_HUMAN_Fowler2018 | 245 | 2643 |
| UBC9_HUMAN_Roth2017 | 158 | 2280 |
| YAP1_HUMAN_Fields2012-singles | 504 | 313 |

**Table S2:** ESM model variants.

| Name | Layers | Parameters | Embedding Dimension | Dataset Used | Tasks | In Zero -Shot? | In Few -Shot? | Ref. |
| --- | --- | --- | --- | --- | --- | --- | --- | --- |
| ESM-2 | 48 | 15B | 5120 | UR50 | General | False | False | [2] |
|  | 36 | 3B | 2560 |  |  | False | False |  |
|  | 33 | 650M | 1280 |  |  | False | **True** |  |
|  | 30 | 150M | 640 |  |  | False | **True** |  |
|  | 12 | 35M | 480 |  |  | False | **True** |  |
|  | 6 | 8M | 320 |  |  | False | **True** |  |
| ESMFold | 48<br>+36 | 690M<br>+3B | - | PDB<br>+UR50 | Folded Structure | False | False | [2] |
| ESM-IF1 | 20 | 124M | 512 | CATH<br>+ UR50 | Inverse Folding | False | False | [28] |
| ESM-1 | 38 | 670M | 1280 | UR50 | General | False | False | [9] |
|  | 12 | 85M | 768 |  |  | False | False |  |
|  | 6 | 43M | 768 |  |  | False | False |  |
| ESM-MSA-1b | 12 | 100M | 768 | UR50<br>+MSA | Variant Prediction | False | False | [29] |
| ESM-1v (x5) | 33 | 650M<br>x5 | 1280 | UR90 | Variant Prediction | **True** | **True**<br>(1 only) | [1] |

## Footnotes

* In this work, the term *function* is used as the mathematical relationship between a protein’s sequence and its quantitative property of interest. It is not intended to refer specifically to enzymatic mechanics, such as kinetics or activity.

* For this work, the terms *protein* and *enzyme* are used interchangeably. Non-enzymatic proteins are out of scope.

* Note that in April 2023, the FAIR ESM research group was shut down. The former team is currently working on a stealth startup for protein prediction.

## Notes

### Competing Interest Statement

The authors have declared no competing interest.

